# A key role of PIEZO2 mechanosensitive ion channel in adipose sensory innervation

**DOI:** 10.1101/2024.11.18.624210

**Authors:** Yu Wang, Yunxiao Zhang, Verina Leung, Saba Heydari Seradj, Utku Sonmez, Rocio Servin-Vences, Darren Lipomi, Li Ye, Ardem Patapoutian

**Affiliations:** Department of Neuroscience, Dorris Neuroscience Center, Scripps Research, San Diego, United States; Howard Hughes Medical Institute, Chevy Chase, United States; Jacobs School of Engineering, UCSD, San Diego, United States

**Author notes:** Correspondence should be addressed to Li Ye or Ardem Patapoutian.

## Abstract

Compared to the well-established functions of sympathetic innervation, the role of sensory afferents in adipose tissues remains less understood. Recent work revealed the anatomical and physiological significance of adipose sensory innervation; however, its molecular underpinning remains unclear. Here, using organ-targeted single-cell RNA sequencing, we identified the mechanoreceptor PIEZO2 as one of the most prevalent receptors in fat-innervating dorsal root ganglia (DRG) neurons. We found that selective PIEZO2 deletion in fat-innervating neurons phenocopied the molecular alternations in adipose tissue caused by DRG ablation. Conversely, a gain-of-function PIEZO2 mutant shifted the adipose phenotypes in the opposite direction. These results indicate that PIEZO2 plays a major role in the sensory regulation of adipose tissues. This discovery opens new avenues for exploring mechanosensation in organs not traditionally considered mechanically active, such as the adipose tissues, and therefore sheds light on the broader significance of mechanosensation in regulating organ function and homeostasis.

## Introduction

Adipose-brain communication is important for whole-body energy homeostasis. Traditionally, the studies of this interaction have been centered on two primary pathways: the brain’s regulation of adipose activity via sympathetic innervation and the ascending signals from adipose tissue to the brain via circulating factors. Recent research, however, has highlighted an important role of afferent sensory innervation of adipose tissues in metabolic regulation^1,2^.

Compared to their sympathetic counterparts, the afferent pathways innervating the adipose tissue remain underexplored. For example, specific ablation of fat-innervating DRG neurons resulted in altered gene expression in adipose tissues, suggesting there is tonic activity within these afferent pathways even under basal conditions^1^. However, the specific stimuli that activate adipose sensory afferents are still largely unknown. Previous studies have shown that leptin receptor (LepR) and Transient receptor potential vanilloid 1 (TRPV1) are expressed in DRG neurons that innervate fat, and extracellular recording showed that the fat nerves can be activated through intra-pad infusion of exogenous leptin, capsaicin, and free fatty acids^3–5^. However, additional genetic evidence is needed to determine whether these act as endogenous stimuli of the sensory terminals under physiological conditions.

To address this knowledge gap, we first determined the molecular identities of fat-innervating DRG neurons. We performed organ-targeted single-cell RNA sequencing to profile the sensory receptors present in fat-innervating DRGs. We identified *Piezo2* as one of the most abundant sensory channels in these neurons. By employing both loss-of-function (LOF) and gain-of- function (GOF) PIEZO2 genetic models, we unveiled an unexpected role for mechanosensation in regulating adipose function, providing fresh insight into the molecular mechanisms of afferent communication between adipose tissue and the brain.

## Results

### *Piezo2* is abundantly expressed in fat-innervating sensory neurons

To determine the molecular identities of fat-innervating DRG neurons, we performed single-cell sequencing of retrogradely labeled DRG neurons from fat (**Figure 1A**). Specifically, we focused on inguinal white adipose tissue (iWAT) and epididymal white adipose tissue (eWAT), as they play important roles in lipid and glucose metabolism and are innervated by non-overlapping neurons from DRGs at similar vertebral levels^1,6^. We injected fluorescent dye-conjugated retrograde tracers into iWAT and eWAT, and collected fluorescently labeled cells and non-labeled cells (which innervate the skin and other organs at similar spinal levels) by fluorescence-activated cell sorting (FACS) for single-cell sequencing (**Figure 1A**). As previously demonstrated^1^, the DRG neurons that innervate iWAT and eWAT did not overlap with each other by FACS (**Figure S1A**). Importantly, we observed a reduction of *Mrgprd* expression, a marker for cutaneous nociceptors^7^, in the iWAT- and eWAT-DRGs comparing to non-labeled cells, consistent with the expectation that our fat-targeted scRNAseq is specifically devoid of skin-innervating neurons. Many known, heterogenous DRG populations^8^ were well-represented in the clustering of fat-innervating DRG neurons (**Figure 1B-C**, S1C). In terms of individual genes, several sensory receptor channels were expressed equally or in higher percentages in iWAT- DRGs compared to unlabeled-DRGs, such as *Piezo2*^9–11^, *Asic1*^12–14^, *Lepr*^4,15^, *Slc17a8*^16–18^, and *Trpm8*^19–21^. Of these, *Piezo2* was the most highly expressed gene (**Figure 1D-E**).

**Figure 1.**
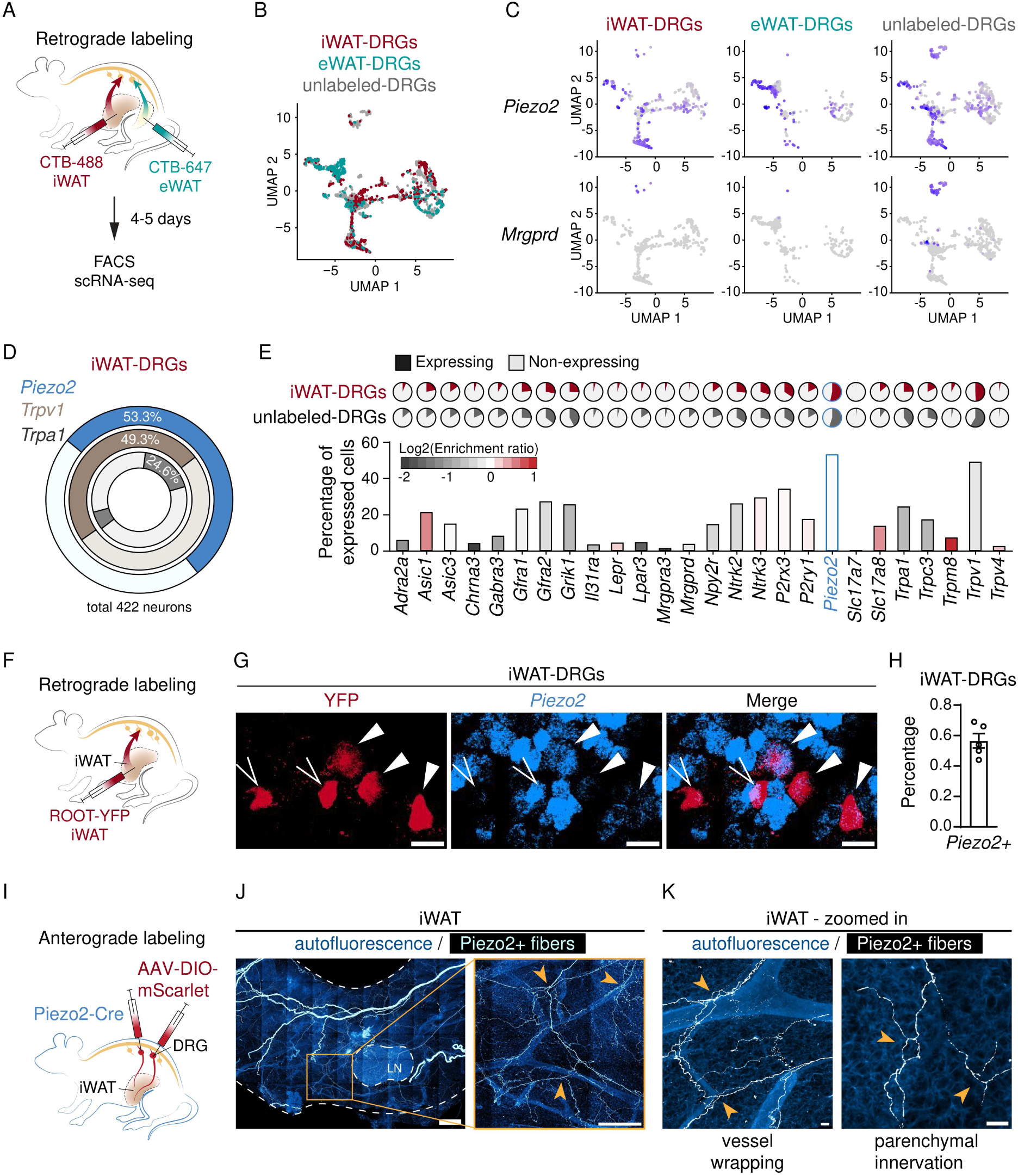
PIEZO2 is expressed in fat-innervating neurons (see also S1) **A.** Schematics of fat-targeted single-cell RNA sequencing. CTB-488 was injected in iWAT, CTB-647 was injected in eWAT, and DRGs from T10-L6 were dissected, dissociated, FACS sorted, and loaded for 10x Genomics scRNA-seq. **B.** UMAP projection of iWAT-DRGs, eWAT-DRGs, and unlabeled-DRGs. **C.** Expression of *Piezo2* and *Mrgprd* in DRG samples. **D.** Quantification of *Piezo2*, *Trpv1*, and *Trpa1* expression percentage in iWAT-DRGs. **E.** Bar plot showing the percentage of indicated genes expressed in iWAT-DRGs and color-coded by the enrichment ratio (defined by the ratio between the percentage in iWAT-DRGs and unlabeled-DRGs). **F.** Schematics of retrograde viral labeling from iWAT. **G.** Representative optical section images of YFP staining and *Piezo2* HCR in whole-mount DRG. **H.** Quantification of *Piezo2* percentage in YFP-positive DRGs, N =5. **I.** Schematics of anterograde viral labeling in PIEZO2-Cre. AAV-DIO-mScarlet was injected into T13/L1 DRGs of PIEZO2-Cre. **J.** Representative images of cleared iWAT tissues after anterograde labeling and zoomed-in views of regions around lymph node. **K.** Zoomed in view of PIEZO2+ sensory nerves in iWAT, showing vessel wrapping and parenchymal innervation morphologies. Scale bar: 30 um in **G, K**, 500 um in **J**.

PIEZO2 is widely recognized as a specialized mechanoreceptor, playing crucial roles in touch^9^, pain^22^, and proprioception^23^. Additionally, it has been increasingly associated with the regulation of internal organ functions, such as baroreception^24^, pulmonary regulation^25^, bladder control^10^, and gastrointestinal activities^11^. However, fat tissue is not conventionally considered an organ that actively receives or responds to mechanical stimuli, in comparison to organs like the lung or gut; therefore, the abundance of PIEZO2 in fat-innervating DRG neurons is striking.

Nevertheless, there have been reports that preadipocytes and adipocytes are mechanosensitive and express sensory channels such as PIEZO1 and SWELL1 that autonomously regulate adipogenesis and lipid accumulation^26,27^, suggesting dynamics in mechanical characteristics of fat is physiologically relevant, and therefore such mechanical information could be transmitted to the CNS through sensory afferents.

To validate the expression of PIEZO2 in fat-innervating neurons beyond scRNA seq, we first examined *Piezo2* transcript levels by in situ hybridization in iWAT-DRGs, owing to their significant implications for human health^28^ and the availability of recently developed and validated tools for studying iWAT-DRGs. Retrograde viral labeling from iWAT using a recently developed AAV capable of retrograde labeling of DRG neurons in a target-specific manner (termed ROOT, or retrograde virus optimized for organ targeting^1^ and hybridization chain reaction against *Piezo2* in whole-mount DRG revealed that approximately half of the iWAT-innervating neuronal soma expressed *Piezo2* (**Figure 1F-H**), echoing 53% in scRNA seq results (**Figure 1E**). To test if the axons of these PIEZO2+ neurons are present in the fat, we injected AAV-expressing Cre-dependent fluorescent protein into the thoracolumbar (thoracic level T13/L1) DRGs of Piezo2-ires-Cre mice^29^ (**Figure 1I**, S1D). HYBRiD-based tissue clearing^30^ and immunolabeling of the iWAT from the injected animals revealed robust PIEZO2+ axons projections into the iWAT. Furthermore, these PIEZO2+ fibers exhibit two predominant morphological features: they travel along the vessels or through the parenchyma in close apposition with adipocytes (**Figure 1J-K**, S1E-F), similar to the morphologies of pan-DRG neurites in fat^1^.

These results collectively demonstrate that PIEZO2 channel is expressed in iWAT-innervating DRGs, leading us to hypothesize a functional role in regulating adipose tissue functions.

#### PIEZO2 deletion leads to DRG ablation-like molecular phenotypes in adipose tissue

To investigate whether PIEZO2 is required for the regulation of adipose tissue, we genetically deleted PIEZO2 in fat-innervating DRG neurons. Specifically, we injected the ROOT AAV expressing Cre-YFP or YFP (as a control on contralateral side) into the iWAT of PIEZO2^fl/fl^ mice^29^ (**Figure 2A**). Based on subsequent in situ hybridization analysis, we deleted PIEZO2 in approximately 60% of iWAT-innervating DRGs (**Figure 2B-C**) in a Cre-dependent manner, comparable to previously reported^1^ DRG ablation efficiency; whereas the total count of labeled fat-innervating neurons remained unchanged (**Figure S2A**). Fat tissues were analyzed 3-4 weeks after the surgery following the same protocol used in previous DRG ablation experiments^1^. This timeframe was chosen to minimize acute complications from the surgery or potential long-term compensations. Notably, compared to the control (Cre-) contralateral sides, the fat pads that received sensory-specific PIEZO2^KO^ (Cre+) were enlarged and exhibited upregulation of genes involved in thermogenesis and de novo lipogenesis (DNL) as well as *Adrb3* (adipocyte-specific adrenergic receptor), mirroring previously reported adipose phenotypes after the DRG-ablation^1^ (**Figure 2D-E**). These changes were observed only in the injected iWAT but not in other distant fat depots such as interscapular brown adipose tissue (iBAT) and eWAT (**Figure S2B, S2D, S2E**), indicating that PIEZO2-mediated effects were specific to the local afferents. Bulk RNA sequencing was then performed to assess the unbiased transcriptomic profiles in the iWAT (**Figure S2C**). Interestingly, the transcriptomic changes induced by PIEZO2^KO^ strongly correlate with those resulting from DRG-ablation (**Figure 2F**), indicating that PIEZO2^KO^ functionally mimics DRG-ablations in terms of regulating adipose functions.

**Figure 2.**
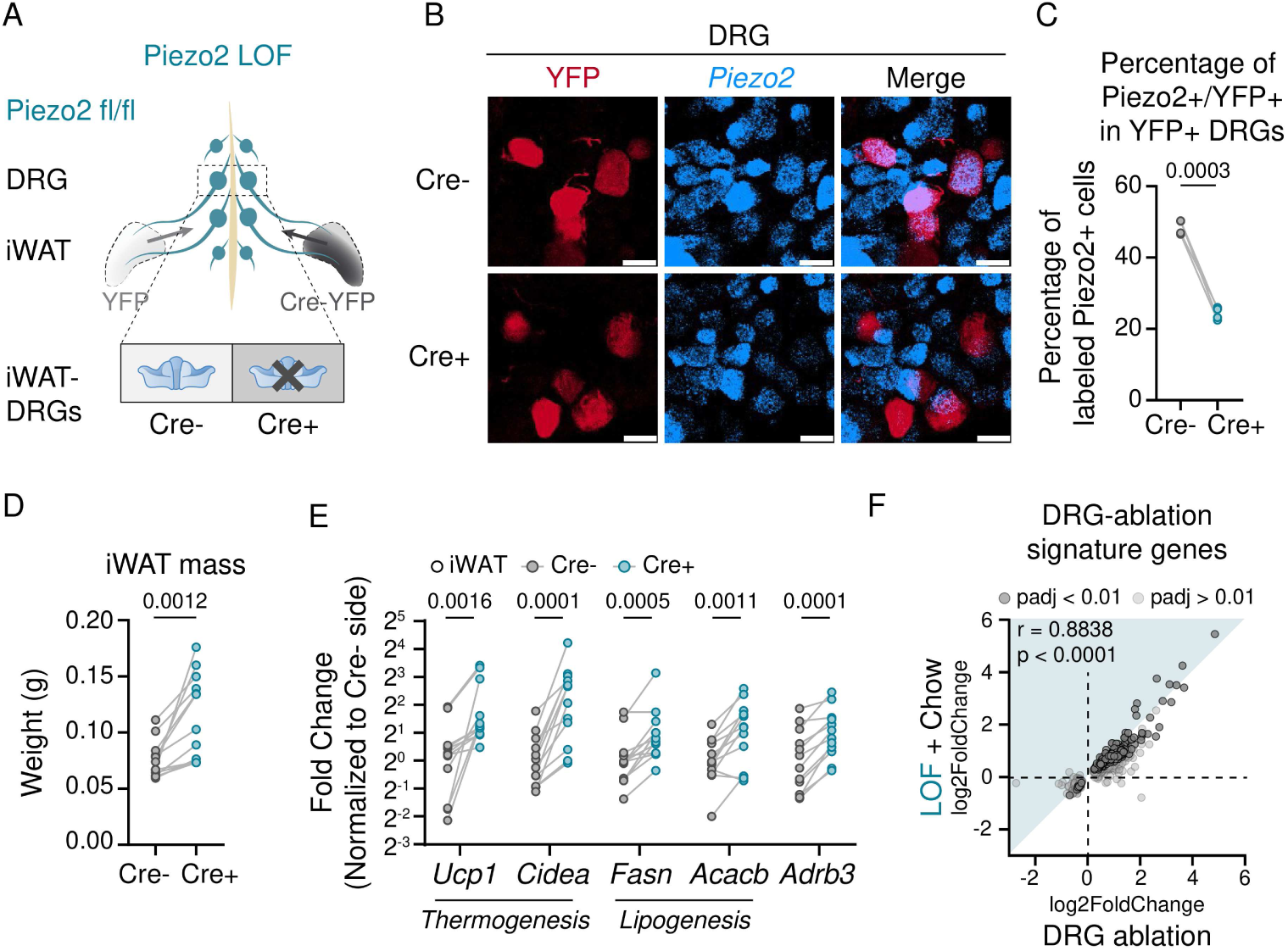
PIEZO2 deletion mimics DRG ablation-induced gene expression changes in fat (see also S2) **A.** Schematics of unilateral fat-DRG PIEZO2 deletion. Retrograde virus expression Cre-YFP or YFP were injected into iWAT of PIEZO2^fl/fl^ mice. **B.** Representative optical section view of *Piezo2* HCR after PIEZO2^KO^. Scale bar: 30 um. **C.** Quantification of Piezo2 knock-out efficiency in iWAT-DRGs. N = 4. Statistics determined by two-tailed paired t-test. **D.** iWAT fat mass changes after PIEZO2^KO^. N = 12. Statistics determined by ratio paired t-tests. **E.** Relative expression level of indicated genes (normalized to Cre-side) in iWAT. N = 12. Statistics determined by multiple paired tests controlling false discovery rate (FDR) by the original FDR method (Benjamini and Hochberg). **F.** Fold change correlation between PIEZO2^KO^ and DRG-ablation (reproduced from Wang et al.^1^) for DRG-ablation signature genes (defined by p_adj_ < 0.01 induced by DRG-ablation). N=6 for PIEZO2^KO^, N=5 for DRG-ablation. Nonparametric Spearman correlation and two-tailed p-value were calculated.

#### PIEZO2 GOF reverses adipose gene expression changes induced by DRG ablation

The similarity between PIEZO2 deletion and DRG ablation on fat gene expression suggests PIEZO2 may play a key role in the fat-innervating sensory afferents. To further test this hypothesis, we sought to investigate if PIEZO2 activity could bi-directionally impact adipose gene expression. We have previously demonstrated that a gain-of-function (GOF) mutation in the C-terminus of Piezo2 (PIEZO2^E2727del^) causes slower channel inactivation kinetics.

Consequently, this mutation elicits larger PIEZO2-dependent currents in response to a given mechanical stimulation, and is associated with distal arthrogryposis in humans^31^. Building on these findings, we generated a transgenic mouse model, which conditionally expresses the analogous mouse PIEZO2 GOF mutant (PIEZO2^E2799del^) in a Cre-dependent manner. This model similarly develops joint contractures, paralleling the human condition^32^.

Using the ROOT-based unilateral targeting strategy, we expressed PIEZO2^E2799del^ mutant in fat- innervating DRGs (**Figure 3A**). With chow diet, we did not observe transcriptional changes in the fat depots between Cre+ and Cre- sides (**Figure 3B**). Given that the characteristic of the GOF mutant lies in the slower closure of the opened channel rather than in overexpression or a lower activation threshold, we hypothesized that a more substantial mechanical trigger may be needed to elicit differential activity between WT and GOF mutant. High fat diet (HFD), which is widely used to model obesity in rodents^33^, can lead to lipid accumulation within adipocyte, ECM remodeling and fat expansion—all of which could potentially increase mechanical stress in adipose tissue^34^. To minimize potential confounding factors associated with late-stage obesity^35–37^, we put mice on HFD for 2-3 weeks, which is sufficient to induce fat expansion without major insulin resistance.

**Figure 3.**
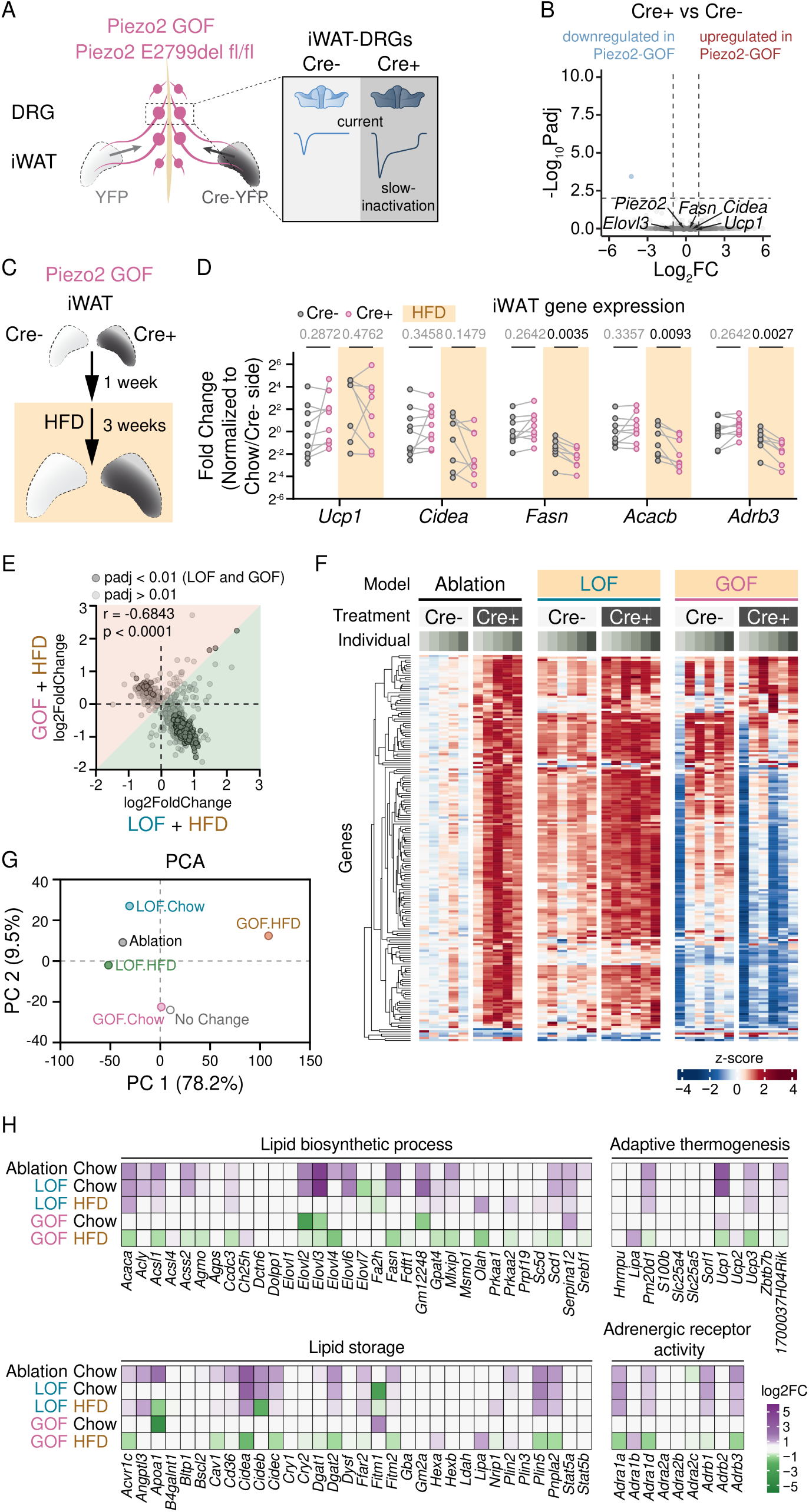
PIEZO2 gain-of-function (GOF) partially reverses molecular changes (see also S3) **A.** Schematics of unilateral fat-DRG PIEZO2^E2799del^ GOF. Retrograde virus expression Cre-YFP or YFP were injected into iWAT of PIEZO2^E2799del^ fl/fl mice. **B.** Volcano plot showing the transcriptional changes in iWAT after PIEZO2^E2799del^ GOF. **C.** Schematics of unilateral fat-DRG PIEZO2^E2799del^ GOF and high-fat diet (HFD) treatment. **D.** Relative expression level of indicated genes (normalized to Chow condition Cre-side) in iWAT under Chow and HFD conditions. N = 9 for chow and N = 8 for HFD. Statistics determined by multiple paired tests controlling false discovery rate (FDR) by the original FDR method (Benjamini and Hochberg). **E.** Fold change correlation GOF-HFD and LOF-HFD for DRG-ablation signature genes. N=6 for each group. Nonparametric Spearman correlation and two-tailed p-value were calculated. **F.** Heatmap showing the normalized expression level in indicated conditions (DRG-ablation, LOF-HFD, GOF-HFD) for top changed genes induced by DRG-ablation. **G.** Principal component analysis of all gene fold changes in all indicated conditions. **H.** Heatmap showing gene fold change for individual genes involved in indicated pathways.

As expected, short-term HFD mildly increased body weight and fat mass (by 2-3 fold) (**Figure S2G-S2J, S3A-S3B**). Also consistent with the literature, HFD downregulated DNL and adrenergic signaling genes, such as *Fasn*, *Acacb* and *Adrb3*, but not thermogenic genes (such as *Ucp1*), potentially due to diet-induced thermogenesis^38–41^ (Figure 3D, S3C-D). Remarkably, under HFD conditions, fat pads from the side expressing the PIEZO2^E2799del^ GOF mutant (i.e., Cre+ side) exhibited gene expression changes in the opposite direction as those induced by PIEZO2 deletion (**Figure 3C-D**, **Figure S3C-D**). Specifically, PIEZO2^E2799del^ GOF mutant further decreased the mRNA levels of *Fasn*, *Acacb* and *Adrb3* compared to the contralateral controls despite the overall expression of these genes being already downregulated by the HFD. In other words, the expression of GOF PIEZO2 in sensory neurons effectively exacerbated the adipose gene expression changes caused by HFD.

Beyond individual genes, global transcriptional analysis of the PIEZO2 KO and GOF iWAT under HFD conditions revealed that PIEZO2^KO^ showed similar upregulation patterns compared to DRG-ablation, while PIEZO2^E2799del^ showed reversed changes compared to the DRG-ablation (and PIEZO2^KO^) (**Figure 3E-3F, S3F-S3G**). Principal component analysis of transcriptomic changes induced by all conditions indicated that PIEZO2^KO^ and DRG-ablation clustered together, while the PIEZO2^E2727del^ and HFD combination shifted the transcriptional profiles to the opposite direction (**Figure 3G**). More specifically, for genes involved in lipid metabolism and adrenergic activity, PIEZO2^E2727del^ showed a regulatory effect opposite to both PIEZO2^KO^ and DRG- ablation (**Figure 3H**) under HFD challenges. Taken together, these results suggest that PIEZO2 activity plays a major role in the afferent regulation of adipose tissue and that HFD-induced fat expansion is likely to further elevate PIEZO2 activity. This mechanical afferent pathway is suppressing sympathetic-related gene programs in the fat, which can be abolished by DRG- ablation and PIEZO2^KO^, but amplified by a slower inactivating mutant of PIEZO2.

## Discussion

Here we describe the role of PIEZO2 in sensory neurons that innervate fat. By organ-targeted scRNA-seq and viral tracing, we found PIEZO2 is among the most abundant receptors expressed in fat-innervating neurons. PIEZO2^KO^ in these neurons largely recapitulates DRG-ablation induced gene changes in iWAT, while expression of a GOF PIEZO2^E2727del^ mutant partially reverses the direction of the changes. These results suggest that mechanosensation plays an unexpected role in regulating fat activity, and raise several questions for future research.

The first question is where are the mechanical stimuli coming from? Typically, the presence of PIEZO1/2 indicates the detection of external forces such as touch and breathing. However, recent studies suggest that these channels can also be activated at the cellular level during normal physiological processes, as demonstrated in red blood cells^42,43^, keratinocytes^44^, and macrophages^45,46^. Although fat has not traditionally been considered a mechanically active organ that experiences rapid volumetric changes, it is highly dynamic and expanding and shrinking in response to metabolic needs^47–49^. Growing evidence suggests that active mechanoactivity occurs within adipose tissues, involving interactions among various cell types. For example, adipocytes and preadipocytes encounter shear stress, compression, and membrane tension. These mechanical forces activate ion channels and signaling pathways, influencing fat cell development and differentiation^26,27,50–55^. Additionally, the ECM in adipose tissues presents physical restraints to fat expansion. In obesity, the ECM becomes stiffer, contributing to pathological alterations such as ectopic fat deposition and insulin resistance^56–62^. Moreover, the mechanical stimuli could also be from vasodilation, and lymphatic pumping, as suggested by the observed close proximity of PIEZO2+ nerves and vasculatures (**Figure S1F**). Together, it is conceivable that one or more mechanical signals from the adipose tissue could reflect the metabolic or energy storage states of this organ which could be transmitted to the brain through the sensory afferents.

However, identifying the specific mechanical stimuli in adipose tissues in vivo is difficult, and limited by available tools. Implantable pressure sensors are designed for organs with either an open lumen or a large stretchable surface, but not for small-sized, solid tissues like the fat.

While small-molecule dye-based tension sensors have been successfully used in vitro, their in vivo application has been complicated by limited tissue permeability, insufficient signal-to-noise ratio, and lack of cellular selectivity. Newer genetically encoded tension or force sensors show promise^63–65^; however, real-time imaging of fine axonal terminals that are deeply embedded in the tissue is yet to be demonstrated. These complexities highlight the need for advanced technological solutions to accurately measure and understand the mechanical stimuli in vivo.

Another question is what else is being sensed by the sensory afferents in the fat? Previous studies suggest that fat-associated nerves can be activated by leptin and lipolysis products^4,5^. In addition, our study has identified many other potential receptors expressed in fat-DRGs through organ- targeted scRNA-seq (**Figure 1E** and S1B). Apart from *Piezo2*, *Asic1* and *Trpm8* were found to be enriched in fat-DRGs comparing to the unlabeled-DRGs, suggesting that acidity or temperature changes could also serve as endogenous stimuli for fat afferents. Additionally, *Trpv1*, which is known to play a crucial role in metabolic homeostasis by whole-body knockout animal models, is also highly expressed in fat-DRGs, and can be activated by inflammatory signals as suggested in other organs such as the lung and the gut^66,67^.

Overall, our findings suggest that PIEZO2 serves as the key mediator for adipose afferent activity. This discovery opens up new directions for investigating mechanosensation in non- traditional mechanoactive organs and provides important insights into mechanosensation in maintaining whole-body homeostasis.

### Limitations of the study

A conceptual limitation is that we have been using PIEZO2 function to infer the involvement of mechanosensation, primarily due to the lack of reliable techniques to directly measure mechanical force at axonal terminals *in vivo*. However, our premise is based on now well- established evidence that PIEZO2 is exclusively activated by mechanical stimuli but not other endogenous activators^23,68–74^. Given the ample precedence in the literature, mechanosensation remains the most likely endogenous signal that activates PIEZO2. Second, when we used ROOT to retrogradely target PIEZO2 gene in the sensory neurons, we could not rule out the possibility that the Cre could also act on local adipose cells to introduce confounding factors. However, considering the expression of PIEZO2 was extremely low and sparse in adipose tissues^27,75^ in comparison to the DRG neurons, it is less likely non-neuronal PIEZO2 would play a major role in the observed phenotype. Furthermore, the striking resemblance between the fat transcriptomic profiles induced by PIEZO2^KO^ and DRG-ablation suggests that afferent PIEZO2 is the main contributor.

## Acknowledgments

We thank all members of the Patapoutian and Ye lab for their support and feedback. We thank Dr Jeffery Friedman, Enrique Saez, Kara Marshall for their input; Jeffery Stirman for the imaging support; the staff at Scripps FACS core and genomics core for sample preparation. This work was supported by the Howard Hughes Medical Institute; NIH grants R35 NS105067 and R01AT012051 (to A.P.); NIH Director’s New Innovator Award DP2DK128800 (to L.Y.), NIDDK K01DK114165 (to L.Y.). L.Y. was also supported by the NIDDK, NIDA, NIMH, NCI, BRAIN Initiative, the Chan Zuckerberg Initiative, and the Dana, Whitehall, Baxter, and Abide-Vividion Foundations, Y.W. was supported by the Dorris Scholar Award, Y.Z. is a Merck Fellow of the Damon Runyon Cancer Research Foundation, DRG-2405-20.

## Author contributions

Y.W., L.Y. and A.P. conceived and designed the study. Y.W., Y.Z., V.H.L., S.H.S., U.S. and M.R.S.-V. performed the experiments and analyzed the data. Y.W., A.P. and L.Y. wrote the manuscript with input from all of the authors.

## STAR METHODS

### RESOURCE AVAILABILITY

#### Lead contact

Inquiries regarding resources, reagents, and further information should be directed to and will be fulfilled by the Lead Contact, Ardem Patapoutian

### Materials availability

This study did not generate new unique reagents. Established mouse lines and other reagents are available upon request from the lead contact.

### Data and code availability

Raw data from single-cell RNAseq and bulk RNA-seq will be deposited at Gene Expression Omnibus (GEO) database and made publicly accessible upon publication. Original code for data analysis will be available on Github and be publicly available upon publication. Any additional information is available from the lead contact upon request.

## EXPERIMENTAL MODEL AND SUBJECT DETAILS

### Animals

Mice were group-housed in standard housing with 12:12h light:dark with ad libitum access to chow diet and water unless specified. The room temperature was kept around 22 °C and humidity between 30-80% (not controlled). For high-fat diet (HFD) feeding experiments, mice were single housed. Mice at least 6 weeks of age from the following strains were used for this study: wild-type (WT) C57BL/6J (Jackson #000664), PIEZO2-Cre (Piezo2^tm1.1(cre)Apat^ ^29^, Jackson #027719), PIEZO2^fl/fl^ (Piezo2^tm2.2Apat^ ^29^, Jackson #027720), PIEZO2 ^E2727del^ ^fl/fl^ (C57BL/6NTac- *Fam38btm3287(ED2799D)Arte* ^32^). PIEZO2^fl/fl^ and PIEZO2 ^E2727del^ ^fl/fl^ were maintained on C57BL/6J background, while PIEZO2-Cre were on CD-1;C57BL/6J background. Both male and female mice were used for anatomical mapping studies, while male mice were used for functional experiments. All animal use protocols were approved by The Scripps Research Institute Institutional Animal Care and Use Committee and were in accordance with the guidelines from the NIH.

### Cell lines

HEK293FT (Invitrogen R70007), used for AAV packaging, were cultured in DMEM (Gibco 11995073) with 1% penicillin/streptomycin (Sigma-Aldrich P4333) and 5% Fetal bovine serum (FBS) (Gibco 10437028) and maintained in incubator at 37°C and 5% CO2.

## METHOD DETAILS

### Adeno-Associated Viruses (AAVs)

PHP.S-EF1a-DIO-mScarlet (capsid, Addgene #103006) and MacPNS.1-EF1a-DIO-mScarlet (capsid^76^ Addgene #185136) were used for anterograde mapping. ROOT-CAG-iCre-YFP, ROOT-CAG-iCre, ROOT-CAG-YFP (capsid, Addgene #192262) were used for retrograde labeling and functional studies. The AAV plasmids were cloned in house, and AAVs were packaged in-house using the published protocol^77^. AAVs were titrated by quantitative PCR, aliquoted into 6-10 μL and flash frozen for long-term storage.

### Surgeries

Mice were anesthetized using isoflurane (4% for induction, 1.5-2% for maintenance). The skin in the surgical area was shaved, hair removed, and the area sterilized using ethanol and iodine.

Post-operatively, mice received a subcutaneous injection of flunixin and topical antibiotic ointment for pain management and infection prevention.

For CTB retrograde labeling, each animal received bilateral injections. Each iWAT and eWAT depot received 4–5 μL of 0.1% CTB-488 (Invitrogen C34775) or CTB-647 (Invitrogen C34778) in PBS, respectively. DRGs were harvested 4-5 days post-surgery for single-cell transcriptomic analysis.

For intraganglionic DRG injection, PHP.S- or MacPNS.1-EF1a-DIO-mScarlet was injected into PIEZO2-Cre mice at 6E13 VG/mL, 200 nL per ganglia following previously described procedure. Tissues were harvested one month post-surgery for tissue clearing and immunostaining.

For ROOT virus injections into fat pads, ROOT AAVs carrying various cargo were injected at 4E13 VG/mL by 2 uL per pad, into iWAT of WT, PIEZO2^fl/fl^, or PIEZO2 ^E2727del^ ^fl/fl^ mice following previous procedures. Tissues were harvested 3-4 weeks post-surgery for weight measurement and transcriptomic analysis.

### Preparation of single-cell suspensions

For DRG single-cell suspension preparation, 16 CTB injected animals and 2 non-injected animals (male, age 7-8 weeks) were used. Animals were sacrificed by isoflurane followed by decapitation. DRGs (T10-L6) were dissected in complete hibernate medium (Hibernate-A medium containing B27 (2%), Glutamax (0.5 mM) and actinomycin D (5 ug/mL, Sigma A1410). Post-dissection, the tissues were briefly spun at 100g at 4°C. The pellet was digested with digestion buffer (Hibernate-A medium (Gibco A1247501), B-27 (2%, Gibco 17504001), Glutamax (0.5mM, Gibco 35050061), Collagenase IV (6.25 mg/mL, Gibco 17104-019), Dispase II (240 U/mL, Roche 04942078001), HCl (6 mM), Actinomycin D (20 ug/mL)) for 30 min at 37°C, followed by further digestion with TrypLE (Gibco 12605010) with Actinomycin D (20 ug/mL) for 5 min at 37°C, which was subsequently quenched with FBS. The cells were then centrifuged at 100g, resuspended in complete hibernate medium, and triturated with 0.5% BSA-coated polished glass pipettes. The cell suspension was loaded on top of Optiprep/complete hibernate medium gradient (455 μL/90 μL) and centrifuged at 100g for 10 min at 4°C. Cell pellets were resuspended in complete hibernate medium for fluorescence-activated cell sorting.

### Fluorescence-activated cell sorting

Fluorescence-activated cell sorting was performed at Scripps FACS core. The resuspended DRG single cell suspension was loaded on a Beckman Coulter, Astrios EQ equipped with 355-nm, 405-nm, 488-nm, 561nm, and 640-nm lasers. System was setup with the 100 µm nozzle tip at 25psi with a piezoelectric-frequency of 45.5 kHz. Instrument alignment was verified using the system’s automated QC. Piezo-electric-frequency and drop delay were determined using IntelliSort II. To ensure high sort efficiency and recovery, the sort mode was set to enrich, and the maximum event rate was limited to 1,000 events per second. The sample path was cleaned and rinsed by running 10% beach for 5 minutes and milli-Q water for 10 minutes. Prior to sorting, sample and sort chambers were cooled to 4° C. DRG suspension from control animals were used to set gate and threshold. DRG suspension from injected animals were sorted into 488+, 647+ and double-negative. The 488+ and 647+ and a small portion of double-negative cells were immediately loaded onto 10x Genomics Chromium chip.

### Tissue clearing and staining

Tissue clearing of iWAT from injected PIEZO2-Cre animals was following the previous published protocol. In brief, iWAT tissues were harvested after trans-cardial perfusion of ice-cold PBS and then 4% PFA (Electron Microscopy Perfusion Fixative, 1224SK). The tissues were then dehydrated and delipidated by THF/25% Quadrol gradient and DCM. Samples were then embedded into A1P4 hydrogel (1% acrylamide, 0.125% Bis, 4% PFA, 0.025% VA-044 (w/v), in 1x PBS), polymerized, and then passively cleared with LiOH-Boric-SDS buffer until samples appeared translucent.

The tissues were washed extensively in PBST (PBS with 0.2% Triton X-100) before immunolabeling. The tissues were incubated in Rabbit anti-RFP antibody (Rockland #600-401- 379, 1:400) in PBST for 5 days at RT and then washed with PBST. This was followed by incubation with Donkey anti-Rabbit-AlexaFlour 647 antibody (Jackson Immuno Research Labs #711-606-152, 1:400) for 5 days at RT, and sequential wash 3x3h in PBST. The samples were refractive index matched using EasyIndex (RI 1.52, LifeCanvas) for confocal and lightsheet microscopy imaging.

### Whole-mount HCR RNA-FISH and immunolabeling in DRG

Whole-mount HCR in DRG were performed following a modified version of the protocol established for whole-mount mouse embryos by Molecular Instruments. In brief, DRGs were harvested immediately after euthanasia and fixed overnight in 4% PFA in PBS at 4°C. The samples were then subjected to dehydration and rehydration through a methanol/PBS- 0.1%Tween20 gradient. The rehydrated samples were permeabilized with PBS-0.1% Triton X- 100 for 1h at RT, followed by an overnight fixation in 4% PFA in PBS. Samples were hybridized with the mouse Piezo2-B1 probe (4nM, Molecular Instruments) and amplified using B1- AlexaFluor647 hairpins following manufacturer’s instructions. Post HCR, samples were washed extensively with 5xSSC-0.1% Tween20, and then with PBS-0.1%Tween 20 before sequential immunolabeling. Samples were blocked with PBS-0.1% Tween20 containing 5% donkey serum for 30min at RT, before incubation with Chicken anti-GFP (Aves Labs #GFP-1020, 1:400) for 1h at RT. After 5x5min washes, the samples were incubated with Donkey anti-Chicken- AlexaFluor488 (Jackson Immuno Research Labs #703-545-155, 1:800) for 30min at RT. The samples were washed 5x5min with PBS-0.1% Tween20, before mounted in RapiClear (Sunjin lab) for subsequent confocal imaging.

### Confocal microscopy

Mounted iWAT samples and DRGs were imaged with Olympus FV3000 confocal microscope using one of the following objectives: 4X, 0.28 NA, air (XLFluor, Olympus); 10X, 0.6 NA, water immersion (XLUMPlanFI, Olympus). Images were acquired with Fluoview (v2.4.1.198).

### Lightsheet microscopy

iWAT samples were mounted using 1% agarose/EasyIndex. Mounted samples were imaged inside the SmartSPIM chamber filled with EasyIndex and sealed with mineral oil on the top. Images were acquired using a 3.6X, 0.2 NA objective (LifeCanvas), with 1.8/1.8/2 um XYZ voxel size. Image acquisition was completed with bilateral illumination along the central plane of symmetry within the sample.

### High-fat-diet feeding

Mice that underwent surgical procedures were allowed a recovery period of one week before being singly housed and fed a HFD (Research Diets D12492) and maintained at RT. The HFD was continued for 2-3 weeks. Body weight of the mice were recorded every week prior to euthanasia and tissue collection.

### 10x Chromium Single Cell 3′ and sequencing

Each sample was processed using the Chromium Next GEM Single Cell 3’ Reagent Kits, v3, single index (10x Genomics). Chromium Single Cell 3’ GEM, Library & Gel Bead Kit v3 (10x Genomics; PN-1000092), the Chromium Chip B Single Cell Kit (10x Genomics; PN- 1000073), and the Chromium i7 Multiplex Kit v2 (10x Genomics; PN-120262).

Briefly, single cells were partitioned into Gel Beads-in-emulsion (GEMS) using the Chromium X instrument (10x Genomics). The presence of poly(dT) primer enabled the creation of full-length cDNA from poly-adenylated mRNA. cDNA was amplified to generate adequate mass for library construction. Sequencing libraries were prepared using end repair, A-tailing, adaptor ligation followed by PCR.

Libraries were pooled and sequenced using the NextSeq 500 (Illumina; High-Output v2.5 flow cell) with 28 cycles for read1, 91 cycles for read2, 8 cycles for i7 and a read depth of 86,052 mean reads per cell. The Cell Ranger pipeline together with the mouse transcriptome mm10 were used to align the reads and to generate the gene-cell matrices.

### RNA extraction

Adipose tissues were dissected between 12-2pm and flash frozen in liquid nitrogen. Total RNA was extracted from frozen tissue using TRIzol (Invitrogen 15596026) and RNeasy Mini Kits (Qiagen 74104) following manufacturer’s instructions.

### Reverse Transcription-PCR

For RT-PCR analysis, total RNA was reverse-transcribed using Maxima H Minus First Strand cDNA Synthesis Kit (Thermo Fisher K1652). The resultant cDNA was mixed with primers (Integrated DNA Technology) and qPCRBio SyGreen Blue Mix (Genesee Scientific 17-507) for RT-qPCR using CFX384 real-time PCR system (Biorad). Normalized mRNA expression was calculated using ΔΔCt method, using *Tbp* (encoding TATA-box-binding protein) mRNA as the reference gene. Statistics was performed on ΔΔCt. Primer sequences (forward and reverse sequence, 5’ → 3’, respectively) are *Tbp* (CCTTGTACCCTTCACCAATGAC and ACAGCCAAGATTCACGGTAGA); *Ucp1* (AGGCTTCCAGTACCATTAGGT and CTGAGTGAGGCAAAGCTGATTT); *Cidea* (ATCACAACTGGCCTGGTTACG and TACTACCCGGTGTCCATTTCT); *Fasn* (GGAGGTGGTGATAGCCGGTAT and TGGGTAATCCATAGAGCCCAG); *Acacb* (CGCTCACCAACAGTAAGGTGG and GCTTGGCAGGGAGTTCCTC); *Adrb3* (AGAAACGGCTCTCTGGCTTTG and TGGTTATGGTCTGTAGTCTCGG).

### RNA-library preparation and bulk RNA sequencing

Total RNA samples were prepared into RNAseq libraries using the NEBNext Ultra II Directional RNA Library Prep Kit for Illumina (New England BioLabs E7760L) following manufacturer’s recommended protocol. Briefly, for each sample 100 ngs total RNA was polyA selected, fragmented and then converted to double stranded cDNA followed by ligation of sequencing adapters. The library were then PCR amplified 13 cycles using barcoded PCR primers, purified and pooled before loading onto an Illumina NextSeq2000 P2 flowcell for 100 base single-end sequencing.

## QUANTIFICATION AND STATISTICAL ANALYSIS

### Imaging analysis

The number of YFP and Piezo2 positive cells were manually quantified in whole-mount DRGs using ImageJ. This process was executed in a treatment-blinded manner to ensure unbiased data collection and analysis.

### scRNA-seq analysis

Seurat was used for single cell RNAseq analysis. Genes expressed in less than three cells were excluded, and cells with less than 200 features were removed. All three samples were merged, and cells with more than 10% mitochondrial-DNA derived gene-expression were filtered out. Post-filtering, the cell-gene matrix was normalized. Feature selection was performed with *FindVariableFeatures* within Seurat. The data was scaled and analyzed by principal component analysis (PCA) over selected features. Uniform manifold approximation and projection (UMAP) was used for visualization. Custom R scripts was used to quantify the percentage of cells expressing specific genes in each sample.

### Bulk RNA-seq analysis

Sequenced reads were aligned to the GRCm39 reference genome (Ensembl version 109), and gene counts were quantified using Salmon. Further analysis was performed in R. Estimated counts were calculated using tximport. Differential gene expression analysis, fold change and p-value were calculated by DESeq2. EnhancedVolcano and ComplexHeatmap were used for visualization. The fold change of all four conditions were normalized and all four conditions were normalized and analyzed by PCA and plotted by factoextra. Specifically, DRG-ablation data was reproduced from Wang et al.^1^

### Study design and statistics

The sample size in this study was not determined using statistical methods, but was guided by prior studies and literature in the field using similar experimental paradigms. No data points were excluded, except for mice with deteriorating health issues post-surgery or during the experiment, or where viral targeting was not achieved as determined by qPCR analysis of the viral construct in the fat tissue.

Data collection was conducted blindly, with post hoc registration to respective condition for unbiased analysis. For all analysis other than sequencing, GraphPad Prism was used with the specified statistical tests. In experiments involving unilateral treatments or differing treatments on either side of the body, the left- or right-side assignment for each animal was randomized. Sample sizes for each experiment are reported in the figure legends. All in vivo experiments were repeated at least twice or combined from at least two independent cohorts, yielding consistent results.

## Supplementary Figure Legends

**Supplementary Figure 1.**
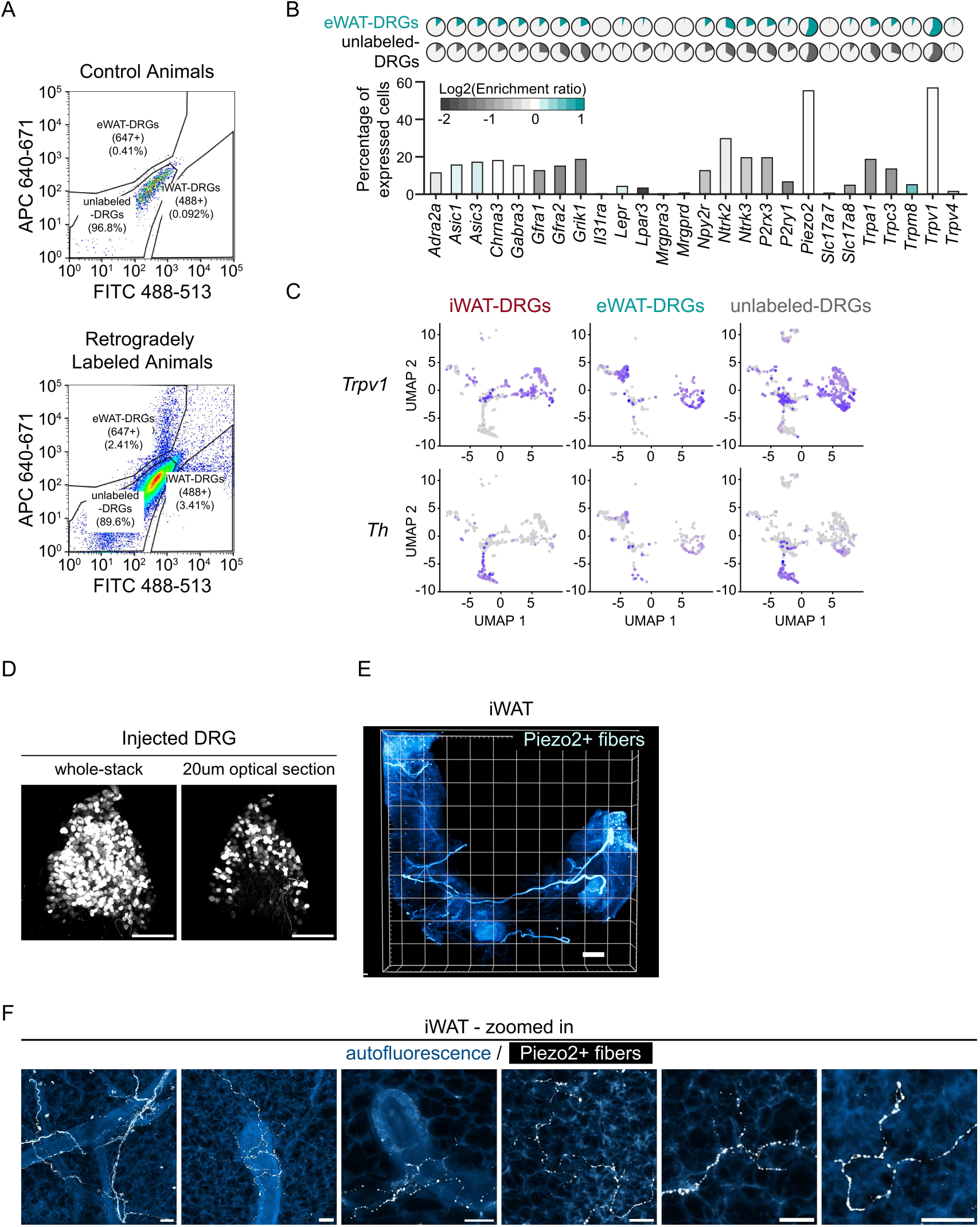
Additional characterization of Piezo2 in fat-innervating DRGs. **A.** Flow cytometric analysis of 488+ (iWAT-DRGs) and 647+ (eWAT-DRGs) from control and CTB-injected mice. **B.** Bar plot showing the percentage of indicated genes expressed in eWAT-DRGs and color-coded by the enrichment ratio (defined by the ratio between the percentage in eWAT-DRGs and unlabeled-DRGs). **C.** Expression of *Trpv1* and *Th* in DRG samples. **D.** Representative whole-mount view of DRG (L1) in PIEZO2-Cre mouse injected with AAV-DIO-mScarlet. **E.** Whole-mount lightsheet microscopy view of iWAT from PIEZO2-Cre mouse injected with AAV-DIO-mScarlet in T13/L1 DRGs. **F.** Zoomed in view of PIEZO2+ sensory nerves in iWAT. Scale bar: 200 um in **D**, 2 mm in **E**, 30 um in **F**.

**Supplementary Figure 2.**
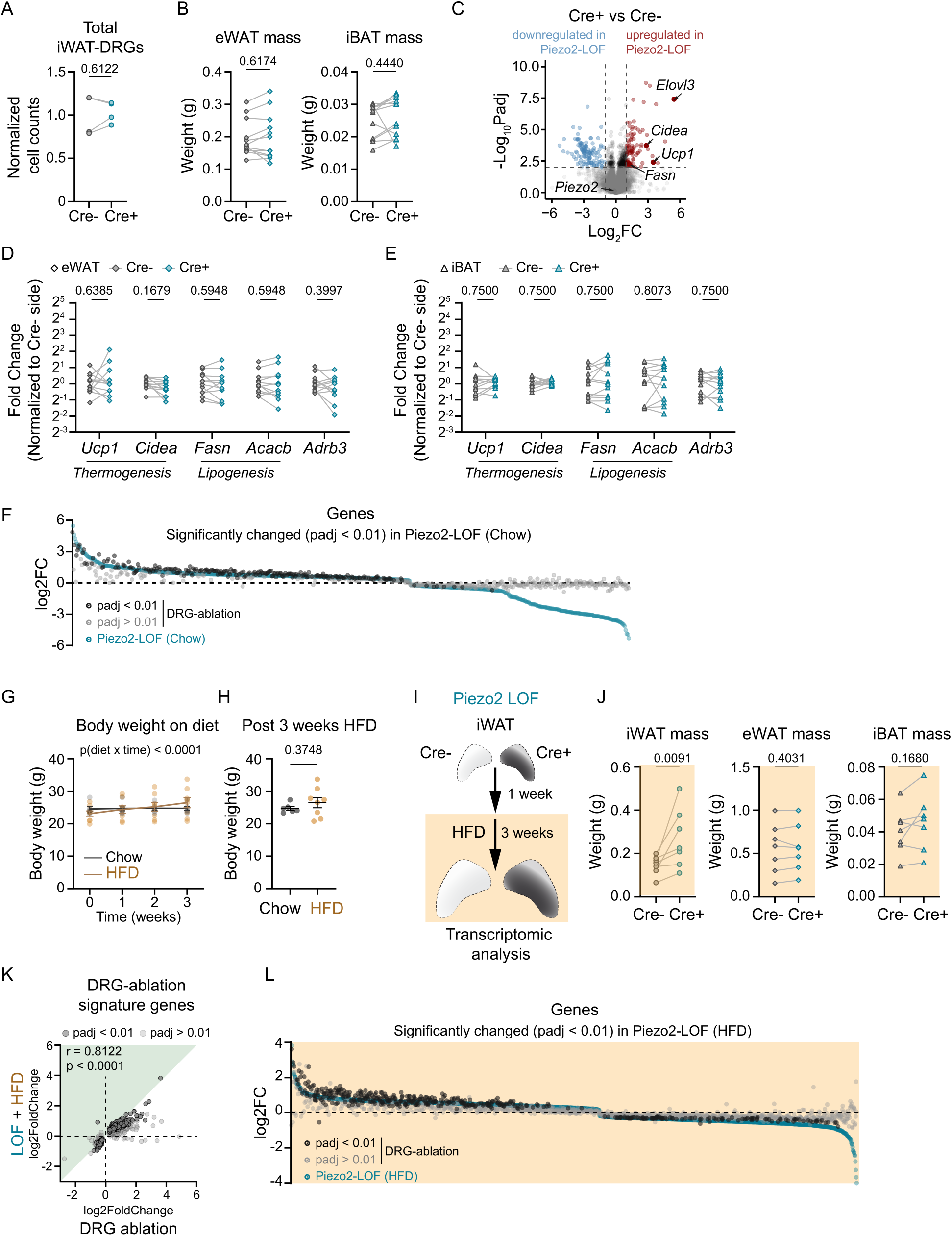
Additional characterization of PIEZO2 LOF. **A.** Normalized total number of retrogradely labeled iWAT-DRGs. N =4. Statistics determined by two-tailed paired t tests. **B.** Fat mass of eWAT and iBAT after fat-DRG PIEZO2^KO^. N = 12. Statistics determined by ratio paired t-tests. **C.** Volcano plot showing gene expression changes in iWAT after PIEZO2^KO^. **D-E**. Relative expression level of indicated genes (normalized to Cre-side) in eWAT (**D**) and iBAT (**E**). N = 12. Statistics determined by multiple paired tests controlling false discovery rate (FDR) by the original FDR method (Benjamini and Hochberg). **F**. Fold change induced by DRG-ablation for all the genes significantly changed by PIEZO2^KO^ (p_adj_ < 0.01). The color shows the p-value. **G-H**. Body weight changes on 3-week high-fat diet. Time-course body weight is shown in (**G**), and body weight after 3 weeks of HFD is shown in (**H**). Statistics were determined by two-way ANOVA in (**G**) and two-tailed t-test (**H**). **I**. Schematics of unilateral PIEZO2^KO^ on HFD. **J**. Fat mass after PIEZO2^KO^ -HFD. N = 7. Statistics determined by ratio paired t-tests. **K**. Fold change correlation between PIEZO2^KO^-HFD and DRG-ablation for DRG-ablation signature genes (defined by p_adj_ < 0.01 induced by DRG-ablation). N = 6 for PIEZO2^KO^-HFD, N = 5 for DRG ablation. Nonparametric Spearman correlation and two-tailed p-value were calculated. **L**. Fold change induced by DRG-ablation for all the genes significantly changed by PIEZO2^KO^-HFD (p_adj_ < 0.01). The color shows the p-value.

**Supplementary Figure 3.**
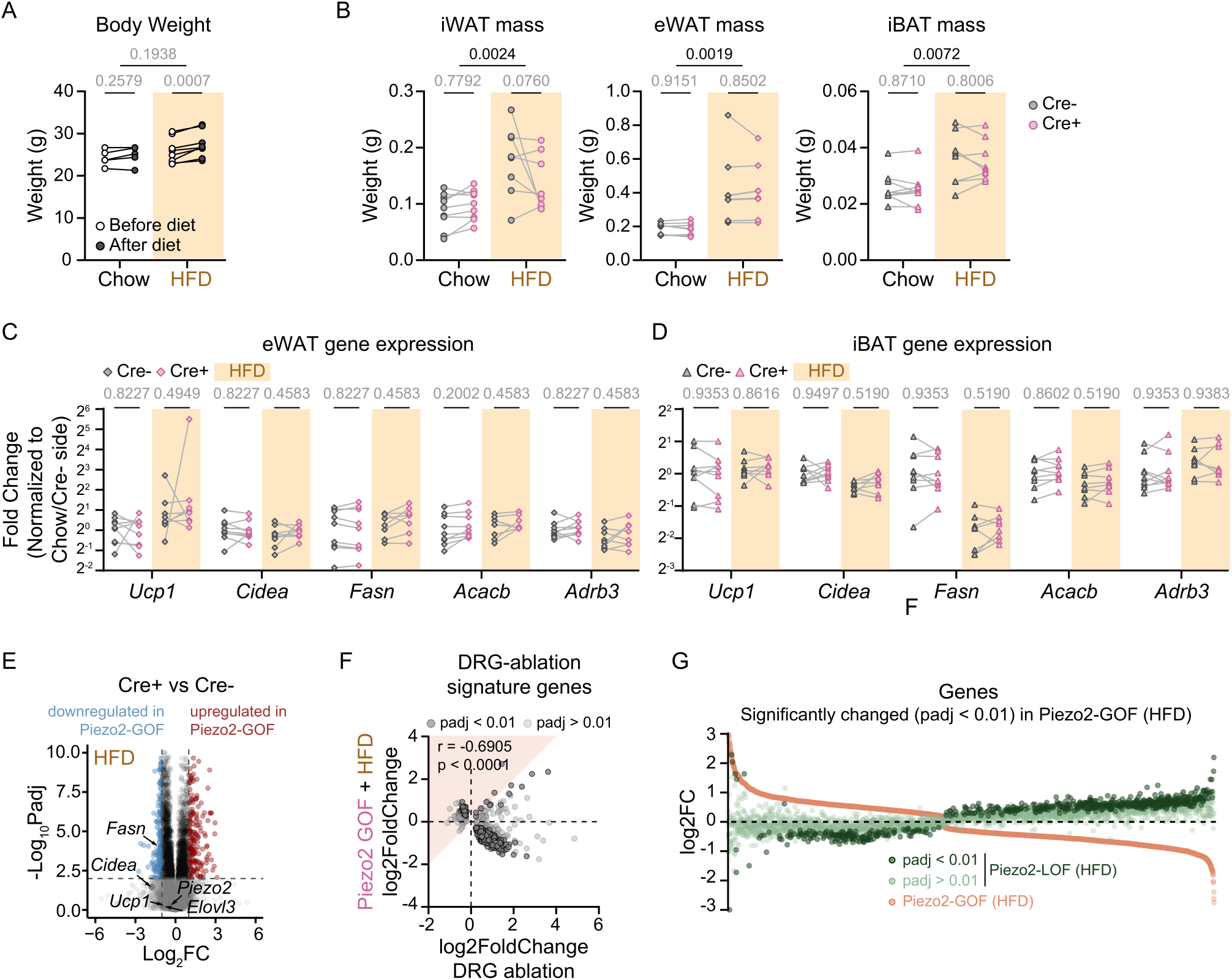
Additional characterization of PIEZO2 GOF. **A**. Body weight changes on 3-week HFD for unilateral PIEZO2^E2799del^ GOF. **B**. Fat mass for PIEZO2^E2799del^ GOF under chow and HFD conditions. N = 9 for GOF-Chow and N = 8 for GOF-HFD. Statistics were determined by two-way ANOVA with Šídák’s multiple comparisons test. **C-D**. Relative expression level of indicated genes (normalized to Chow condition Cre-side) in eWAT and iBAT under Chow and HFD conditions. N = 9 for chow and N = 8 for HFD. Statistics determined by multiple paired tests controlling false discovery rate (FDR) by the original FDR method (Benjamini and Hochberg). **E**. Volcano plot showing transcriptional changes in iWAT after Piezo2 GOF. **F**. Fold change correlation between GOF-HFD and DRG -ablation for DRG-ablation signature genes. N = 6 for GOF-HFD, N = 5 for ablation. Nonparametric Spearman correlation and two- tailed p-value were calculated. **G**. Fold change induced by LOF-HFD for all genes significantly changed by GOF-HFD (p_adj_ < 0.01). The color shows the p-value.

## Notes

### Competing Interest Statement

The authors have declared no competing interest.

## References

1. Wang, Y., Leung, V.H., Zhang, Y., Nudell, V.S., Loud, M., Servin-Vences, M.R., Yang, D., Wang, K., Moya-Garzon, M.D., Li, V.L., et al. (2022). The role of somatosensory innervation of adipose tissues. Nature 609, 569–574.

2. Mishra, G., and Townsend, K.L. (2023). The metabolic and functional roles of sensory nerves in adipose tissues. Nat. Metab. 5, 1461–1474.

3. Shi, Z., Chen, W.-W., Xiong, X.-Q., Han, Y., Zhou, Y.-B., Zhang, F., Gao, X.-Y., and Zhu, G.-Q. (2012). Sympathetic activation by chemical stimulation of white adipose tissues in rats. J. Appl. Physiol. 112, 1008–1014.

4. Murphy, K.T., Schwartz, G.J., Nguyen, N.L.T., Mendez, J.M., Ryu, V., and Bartness, T.J. (2013). Leptin-sensitive sensory nerves innervate white fat. Am. J. Physiol. Endocrinol. Metab. 304, E1338–47.

5. Garretson, J.T., Szymanski, L.A., Schwartz, G.J., Xue, B., Ryu, V., and Bartness, T.J. (2016). Lipolysis sensation by white fat afferent nerves triggers brown fat thermogenesis. Mol. Metab. 5, 626–634.

6. Rosen, E.D., and Spiegelman, B.M. (2014). What we talk about when we talk about fat. Cell 156, 20–44.

7. Dussor, G., Zylka, M.J., Anderson, D.J., and McCleskey, E.W. (2008). Cutaneous sensory neurons expressing the Mrgprd receptor sense extracellular ATP and are putative nociceptors. J. Neurophysiol. 99, 1581–1589.

8. Sharma, N., Flaherty, K., Lezgiyeva, K., Wagner, D.E., Klein, A.M., and Ginty, D.D. (2020). The emergence of transcriptional identity in somatosensory neurons. Nature 577, 392–398.

9. Ranade, S.S., Woo, S.-H., Dubin, A.E., Moshourab, R.A., Wetzel, C., Petrus, M., Mathur, J., Bégay, V., Coste, B., Mainquist, J., et al. (2014). Piezo2 is the major transducer of mechanical forces for touch sensation in mice. Nature 516, 121–125.

10. Marshall, K.L., Saade, D., Ghitani, N., Coombs, A.M., Szczot, M., Keller, J., Ogata, T., Daou, I., Stowers, L.T., Bönnemann, C.G., et al. (2020). PIEZO2 in sensory neurons and urothelial cells coordinates urination. Nature 588, 290–295.

11. Servin-Vences, M.R., Lam, R.M., Koolen, A., Wang, Y., Saade, D.N., Loud, M., Kacmaz, H., Frausto, S., Zhang, Y., Beyder, A., et al. (2023). PIEZO2 in somatosensory neurons controls gastrointestinal transit. Cell 186, 3386–3399.e15.

12. Page, A.J., Brierley, S.M., Martin, C.M., Price, M.P., Symonds, E., Butler, R., Wemmie, J.A., and Blackshaw, L.A. (2005). Different contributions of ASIC channels 1a, 2, and 3 in gastrointestinal mechanosensory function. Gut *54*, 1408–1415.

13. Poirot, O., Berta, T., Decosterd, I., and Kellenberger, S. (2006). Distinct ASIC currents are expressed in rat putative nociceptors and are modulated by nerve injury. J. Physiol. 576, 215–234.

14. Hughes, P.A., Brierley, S.M., Young, R.L., and Blackshaw, L.A. (2007). Localization and comparative analysis of acid-sensing ion channel (ASIC1, 2, and 3) mRNA expression in mouse colonic sensory neurons within thoracolumbar dorsal root ganglia. J. Comp. Neurol. *500*, 863–875.

15. Chen, H.P., Fan, J., and Cui, S. (2006). Detection and estrogen regulation of leptin receptor expression in rat dorsal root ganglion. Histochem. Cell Biol. 126, 363–369.

16. Seal, R.P., Wang, X., Guan, Y., Raja, S.N., Woodbury, C.J., Basbaum, A.I., and Edwards, R.H. (2009). Injury-induced mechanical hypersensitivity requires C-low threshold mechanoreceptors. Nature 462, 651–655.

17. Lou, S., Duan, B., Vong, L., Lowell, B.B., and Ma, Q. (2013). Runx1 controls terminal morphology and mechanosensitivity of VGLUT3-expressing C-mechanoreceptors. J. Neurosci. 33, 870–882.

18. Draxler, P., Honsek, S.D., Forsthuber, L., Hadschieff, V., and Sandkühler, J. (2014). VGluT3^+^ primary afferents play distinct roles in mechanical and cold hypersensitivity depending on pain etiology. J. Neurosci. 34, 12015–12028.

19. Kobayashi, K., Fukuoka, T., Obata, K., Yamanaka, H., Dai, Y., Tokunaga, A., and Noguchi, K. (2005). Distinct expression of TRPM8, TRPA1, and TRPV1 mRNAs in rat primary afferent neurons with adelta/c-fibers and colocalization with trk receptors. J. Comp. Neurol. *493*, 596–606.

20. Dhaka, A., Earley, T.J., Watson, J., and Patapoutian, A. (2008). Visualizing cold spots: TRPM8-expressing sensory neurons and their projections. J. Neurosci. 28, 566–575.

21. Harrington, A.M., Hughes, P.A., Martin, C.M., Yang, J., Castro, J., Isaacs, N.J., Blackshaw, A.L., and Brierley, S.M. (2011). A novel role for TRPM8 in visceral afferent function. Pain 152, 1459–1468.

22. Murthy, S.E., Loud, M.C., Daou, I., Marshall, K.L., Schwaller, F., Kühnemund, J., Francisco, A.G., Keenan, W.T., Dubin, A.E., Lewin, G.R., et al. (2018). The mechanosensitive ion channel Piezo2 mediates sensitivity to mechanical pain in mice. Sci. Transl. Med. 10, eaat9897.

23. Szczot, M., Nickolls, A.R., Lam, R.M., and Chesler, A.T. (2021). The Form and Function of PIEZO2. Annu. Rev. Biochem. 90, 507–534.

24. Zeng, W.-Z., Marshall, K.L., Min, S., Daou, I., Chapleau, M.W., Abboud, F.M., Liberles, S.D., and Patapoutian, A. (2018). PIEZOs mediate neuronal sensing of blood pressure and the baroreceptor reflex. Science 362, 464–467.

25. Nonomura, K., Woo, S.-H., Chang, R.B., Gillich, A., Qiu, Z., Francisco, A.G., Ranade, S.S., Liberles, S.D., and Patapoutian, A. (2017). Piezo2 senses airway stretch and mediates lung inflation-induced apnoea. Nature 541, 176–181.

26. Zhang, Y., Xie, L., Gunasekar, S.K., Tong, D., Mishra, A., Gibson, W.J., Wang, C., Fidler, T., Marthaler, B., Klingelhutz, A., et al. (2017). SWELL1 is a regulator of adipocyte size, insulin signalling and glucose homeostasis. Nat. Cell Biol. 19, 504–517.

27. Wang, S., Cao, S., Arhatte, M., Li, D., Shi, Y., Kurz, S., Hu, J., Wang, L., Shao, J., Atzberger, A., et al. (2020). Adipocyte Piezo1 mediates obesogenic adipogenesis through the FGF1/FGFR1 signaling pathway in mice. Nat. Commun. 11, 2303.

28. Wu, J., Boström, P., Sparks, L.M., Ye, L., Choi, J.H., Giang, A.-H., Khandekar, M., Virtanen, K.A., Nuutila, P., Schaart, G., et al. (2012). Beige adipocytes are a distinct type of thermogenic fat cell in mouse and human. Cell 150, 366–376.

29. Woo, S.-H., Ranade, S., Weyer, A.D., Dubin, A.E., Baba, Y., Qiu, Z., Petrus, M., Miyamoto, T., Reddy, K., Lumpkin, E.A., et al. (2014). Piezo2 is required for Merkel-cell mechanotransduction. Nature 509, 622–626.

30. Nudell, V., Wang, Y., Pang, Z., Lal, N.K., Huang, M., Shaabani, N., Kanim, W., Teijaro, J., Maximov, A., and Ye, L. (2022). HYBRiD: hydrogel-reinforced DISCO for clearing mammalian bodies. Nat. Methods 19, 479–485.

31. Coste, B., Houge, G., Murray, M.F., Stitziel, N., Bandell, M., Giovanni, M.A., Philippakis, A., Hoischen, A., Riemer, G., Steen, U., et al. (2013). Gain-of-function mutations in the mechanically activated ion channel PIEZO2 cause a subtype of Distal Arthrogryposis. Proc. Natl. Acad. Sci. U. S. A. 110, 4667–4672.

32. Ma, S., Dubin, A.E., Romero, L.O., Loud, M., Salazar, A., Chu, S., Klier, N., Masri, S., Zhang, Y., Wang, Y., et al. (2023). Excessive mechanotransduction in sensory neurons causes joint contractures. Science 379, 201–206.

33. Hariri, N., and Thibault, L. (2010). High-fat diet-induced obesity in animal models. Nutr. Res. Rev. 23, 270–299.

34. Lecoutre, S., Lambert, M., Drygalski, K., Dugail, I., Maqdasy, S., Hautefeuille, M., and Clément, K. (2022). Importance of the microenvironment and mechanosensing in adipose tissue biology. Cells 11, 2310.

35. Kwon, E.-Y., Shin, S.-K., Cho, Y.-Y., Jung, U.J., Kim, E., Park, T., Park, J.H.Y., Yun, J.W., McGregor, R.A., Park, Y.B., et al. (2012). Time-course microarrays reveal early activation of the immune transcriptome and adipokine dysregulation leads to fibrosis in visceral adipose depots during diet-induced obesity. BMC Genomics 13, 450.

36. Williams, L.M., Campbell, F.M., Drew, J.E., Koch, C., Hoggard, N., Rees, W.D., Kamolrat, T., Thi Ngo, H., Steffensen, I.-L., Gray, S.R., et al. (2014). The development of diet-induced obesity and glucose intolerance in C57BL/6 mice on a high-fat diet consists of distinct phases. PLoS One 9, e106159.

37. He, M.-Q., Wang, J.-Y., Wang, Y., Sui, J., Zhang, M., Ding, X., Zhao, Y., Chen, Z.-Y., Ren, X.-X., and Shi, B.-Y. (2020). High-fat diet-induced adipose tissue expansion occurs prior to insulin resistance in C57BL/6J mice. Chronic Dis. Transl. Med. 6, 198–207.

38. Jiang, L., Wang, Q., Yu, Y., Zhao, F., Huang, P., Zeng, R., Qi, R.Z., Li, W., and Liu, Y. (2009). Leptin contributes to the adaptive responses of mice to high-fat diet intake through suppressing the lipogenic pathway. PLoS One 4, e6884.

39. Duarte, J.A.G., Carvalho, F., Pearson, M., Horton, J.D., Browning, J.D., Jones, J.G., and Burgess, S.C. (2014). A high-fat diet suppresses de novo lipogenesis and desaturation but not elongation and triglyceride synthesis in mice. J. Lipid Res. 55, 2541–2553.

40. Choi, M.-S., Kim, Y.-J., Kwon, E.-Y., Ryoo, J.Y., Kim, S.R., and Jung, U.J. (2015). High-fat diet decreases energy expenditure and expression of genes controlling lipid metabolism, mitochondrial function and skeletal system development in the adipose tissue, along with increased expression of extracellular matrix remodelling- and inflammation-related genes. Br. J. Nutr. 113, 867–877.

41. Fromme, T., and Klingenspor, M. (2011). Uncoupling protein 1 expression and high-fat diets. Am. J. Physiol. Regul. Integr. Comp. Physiol. 300, R1–8.

42. Cahalan, S.M., Lukacs, V., Ranade, S.S., Chien, S., Bandell, M., and Patapoutian, A. (2015). Piezo1 links mechanical forces to red blood cell volume. Elife 4, e07370.

43. Ma, S., Cahalan, S., LaMonte, G., Grubaugh, N.D., Zeng, W., Murthy, S.E., Paytas, E., Gamini, R., Lukacs, V., Whitwam, T., et al. (2018). Common PIEZO1 allele in African populations causes RBC dehydration and attenuates Plasmodium infection. Cell 173, 443–455.e12.

44. Holt, J.R., Zeng, W.-Z., Evans, E.L., Woo, S.-H., Ma, S., Abuwarda, H., Loud, M., Patapoutian, A., and Pathak, M.M. (2021). Spatiotemporal dynamics of PIEZO1 localization controls keratinocyte migration during wound healing. Elife 10. 10.7554/eLife.65415.

45. Atcha, H., Jairaman, A., Holt, J.R., Meli, V.S., Nagalla, R.R., Veerasubramanian, P.K., Brumm, K.T., Lim, H.E., Othy, S., Cahalan, M.D., et al. (2021). Mechanically activated ion channel Piezo1 modulates macrophage polarization and stiffness sensing. Nat. Commun. 12, 3256.

46. Ma, S., Dubin, A.E., Zhang, Y., Mousavi, S.A.R., Wang, Y., Coombs, A.M., Loud, M., Andolfo, I., and Patapoutian, A. (2021). A role of PIEZO1 in iron metabolism in mice and humans. Cell 184, 969–982.e13.

47. Rutkowski, J.M., Stern, J.H., and Scherer, P.E. (2015). The cell biology of fat expansion. J. Cell Biol. 208, 501–512.

48. Tang, H.-N., Tang, C.-Y., Man, X.-F., Tan, S.-W., Guo, Y., Tang, J., Zhou, C.-L., and Zhou, H.-D. (2017). Plasticity of adipose tissue in response to fasting and refeeding in male mice. Nutr. Metab. (Lond.) 14, 3.

49. Chouchani, E.T., and Kajimura, S. (2019). Metabolic adaptation and maladaptation in adipose tissue. Nat. Metab. 1, 189–200.

50. Hara, Y., Wakino, S., Tanabe, Y., Saito, M., Tokuyama, H., Washida, N., Tatematsu, S., Yoshioka, K., Homma, K., Hasegawa, K., et al. (2011). Rho and Rho-kinase activity in adipocytes contributes to a vicious cycle in obesity that may involve mechanical stretch. Sci. Signal. 4, ra3.

51. Shoham, N., Gottlieb, R., Sharabani-Yosef, O., Zaretsky, U., Benayahu, D., and Gefen, A. (2012). Static mechanical stretching accelerates lipid production in 3T3-L1 adipocytes by activating the MEK signaling pathway. Am. J. Physiol. Cell Physiol. 302, C429–41.

52. El Ouarrat, D., Isaac, R., Lee, Y.S., Oh, D.Y., Wollam, J., Lackey, D., Riopel, M., Bandyopadhyay, G., Seo, J.B., Sampath-Kumar, R., et al. (2020). TAZ is a negative regulator of PPARγ activity in adipocytes and TAZ deletion improves insulin sensitivity and glucose tolerance. Cell Metab. 31, 162–173.e5.

53. Wang, L., Wang, S., Shi, Y., Li, R., Günther, S., Ong, Y.T., Potente, M., Yuan, Z., Liu, E., and Offermanns, S. (2020). YAP and TAZ protect against white adipocyte cell death during obesity. Nat. Commun. 11, 5455.

54. Pope, B.D., Warren, C.R., Parker, K.K., and Cowan, C.A. (2016). Microenvironmental control of adipocyte fate and function. Trends Cell Biol. 26, 745–755.

55. Tharp, K.M., Kang, M.S., Timblin, G.A., Dempersmier, J., Dempsey, G.E., Zushin, P.-J.H., Benavides, J., Choi, C., Li, C.X., Jha, A.K., et al. (2018). Actomyosin-mediated tension orchestrates uncoupled respiration in adipose tissues. Cell Metab. 27, 602–615.e4.

56. 56. Pellegrinelli, V., Heuvingh, J., du Roure, O., Rouault, C., Devulder, A., Klein, C., Lacasa, M., Clément, E., Lacasa, D., and Clément, K. (2014). Human adipocyte function is impacted by mechanical cues: Human adipocytes as mechanosensitive cells. J. Pathol. 233, 183–195.

57. Naftaly, A., Kislev, N., Izgilov, R., Adler, R., Silber, M., Shalgi, R., and Benayahu, D. (2022). Nutrition alters the stiffness of adipose tissue and cell signaling. Int. J. Mol. Sci. 23. 10.3390/ijms232315237.

58. Romani, P., Valcarcel-Jimenez, L., Frezza, C., and Dupont, S. (2021). Crosstalk between mechanotransduction and metabolism. Nat. Rev. Mol. Cell Biol. 22, 22–38.

59. Sun, K., Li, X., and Scherer, P.E. (2023). Extracellular matrix (ECM) and fibrosis in adipose tissue: Overview and perspectives. Compr. Physiol. 13, 4387–4407.

60. Chun, T.-H., Hotary, K.B., Sabeh, F., Saltiel, A.R., Allen, E.D., and Weiss, S.J. (2006). A pericellular collagenase directs the 3-dimensional development of white adipose tissue. Cell 125, 577–591.

61. Khan, T., Muise, E.S., Iyengar, P., Wang, Z.V., Chandalia, M., Abate, N., Zhang, B.B., Bonaldo, P., Chua, S., and Scherer, P.E. (2009). Metabolic dysregulation and adipose tissue fibrosis: Role of collagen VI. Mol. Cell. Biol. 29, 1575–1591.

62. Sun, K., Tordjman, J., Clément, K., and Scherer, P.E. (2013). Fibrosis and adipose tissue dysfunction. Cell Metab. 18, 470–477.

63. Fischer, L.S., Rangarajan, S., Sadhanasatish, T., and Grashoff, C. (2021). Molecular force measurement with tension sensors. Annu. Rev. Biophys. 50, 595–616.

64. Yaganoglu, S., Kalyviotis, K., Vagena-Pantoula, C., Jülich, D., Gaub, B.M., Welling, M., Lopes, T., Lachowski, D., Tang, S.S., Del Rio Hernandez, A., et al. (2023). Highly specific and non-invasive imaging of Piezo1-dependent activity across scales using GenEPi. Nat. Commun. 14, 4352.

65. Sloas, D.C., Tran, J.C., Marzilli, A.M., and Ngo, J.T. (2023). Tension-tuned receptors for synthetic mechanotransduction and intercellular force detection. Nat. Biotechnol. 41, 1287– 1295.

66. Baral, P., Umans, B.D., Li, L., Wallrapp, A., Bist, M., Kirschbaum, T., Wei, Y., Zhou, Y., Kuchroo, V.K., Burkett, P.R., et al. (2018). Nociceptor sensory neurons suppress neutrophil and γδ T cell responses in bacterial lung infections and lethal pneumonia. Nat. Med. 24, 417–426.

67. Lai, N.Y., Musser, M.A., Pinho-Ribeiro, F.A., Baral, P., Jacobson, A., Ma, P., Potts, D.E., Chen, Z., Paik, D., Soualhi, S., et al. (2020). Gut-Innervating Nociceptor Neurons Regulate Peyer’s Patch Microfold Cells and SFB Levels to Mediate Salmonella Host Defense. Cell 180, 33–49.e22.

68. Coste, B., Mathur, J., Schmidt, M., Earley, T.J., Ranade, S., Petrus, M.J., Dubin, A.E., and Patapoutian, A. (2010). Piezo1 and Piezo2 are essential components of distinct mechanically activated cation channels. Science 330, 55–60.

69. 69. Woo, S.-H., Lukacs, V., de Nooij, J.C., Zaytseva, D., Criddle, C.R., Francisco, A., Jessell, T.M., Wilkinson, K.A., and Patapoutian, A. (2015). Piezo2 is the principal mechanotransduction channel for proprioception. Nat. Neurosci. 18, 1756–1762.

70. Chesler, A.T., Szczot, M., Bharucha-Goebel, D., Čeko, M., Donkervoort, S., Laubacher, C., Hayes, L.H., Alter, K., Zampieri, C., Stanley, C., et al. (2016). The role of PIEZO2 in human mechanosensation. N. Engl. J. Med. 375, 1355–1364.

71. Murthy, S.E., Dubin, A.E., and Patapoutian, A. (2017). Piezos thrive under pressure: mechanically activated ion channels in health and disease. Nat. Rev. Mol. Cell Biol. 18, 771–783.

72. Kefauver, J.M., Ward, A.B., and Patapoutian, A. (2020). Discoveries in structure and physiology of mechanically activated ion channels. Nature 587, 567–576.

73. Delmas, P., Parpaite, T., and Coste, B. (2022). PIEZO channels and newcomers in the mammalian mechanosensitive ion channel family. Neuron 110, 2713–2727.

74. Nickolls, A.R., O’Brien, G.S., Shnayder, S., Zhang, Y., Nagel, M., Patapoutian, A., and Chesler, A.T. (2022). Reevaluation of Piezo1 as a gut RNA sensor. Elife 11. 10.7554/eLife.83346.

75. 75. Emont, M.P., Jacobs, C., Essene, A.L., Pant, D., Tenen, D., Colleluori, G., Di Vincenzo, A., Jørgensen, A.M., Dashti, H., Stefek, A., et al. (2022). A single-cell atlas of human and mouse white adipose tissue. Nature 603, 926–933.

76. Chen, X., Ravindra Kumar, S., Adams, C.D., Yang, D., Wang, T., Wolfe, D.A., Arokiaraj, C.M., Ngo, V., Campos, L.J., Griffiths, J.A., et al. (2022). Engineered AAVs for non-invasive gene delivery to rodent and non-human primate nervous systems. Neuron 110, 2242–2257.e6.

77. Challis, R.C., Ravindra Kumar, S., Chan, K.Y., Challis, C., Beadle, K., Jang, M.J., Kim, H.M., Rajendran, P.S., Tompkins, J.D., Shivkumar, K., et al. (2019). Systemic AAV vectors for widespread and targeted gene delivery in rodents. Nat. Protoc. 14, 379–414.

